# Influence of warming temperatures on coregonine embryogenesis within and among species

**DOI:** 10.1101/2021.02.13.431107

**Authors:** Taylor R. Stewart, Mikko Mäkinen, Chloé Goulon, Jean Guillard, Timo J. Marjomäki, Emilien Lasne, Juha Karjalainen, Jason D. Stockwell

## Abstract

The greatest known global response of lakes to climate change has been an increase in water temperatures. The responses of many lake fishes to warming water temperatures are projected to be inadequate to counter the speed and magnitude of climate change. We experimentally evaluated the responses of embryos from a group of cold, stenothermic fishes (Salmonidae Coregoninae) to increased incubation temperatures. Study groups included cisco (*Coregonus artedi*) from lakes Superior and Ontario (USA), and vendace (*C. albula*) and European whitefish (*C. lavaretus*) from Lake Southern Konnevesi (Finland). Embryos from artificial crossings were incubated at water temperatures of 2.0, 4.5, 7.0, and 9.0°C, and their responses were quantified for developmental and morphological traits. Embryo survival, incubation period, and length-at-hatch were inversely related to incubation temperature whereas yolk-sac volume increased with incubation temperature within study groups. However, varying magnitudes of responses among study groups suggested differential levels of developmental plasticity to climate change. Differential levels of parental effects indicate genetic diversity may enable all study groups to adapt to cope with some degree of changing environmental conditions. Our results suggest that the coregonines sampled within and among systems may have a wide range of embryo responses to warming incubation conditions.

## INTRODUCTION

Freshwater lakes are sensitive to climate change (Jenny et al., 2020; Woolway et al., 2020), and one of the greatest threats to lakes from climate change on a global scale is rising water temperatures (Austin & Colman, 2007; O’Reilly et al., 2015; Woolway et al., 2017; Maberly et al., 2020). The greatest seasonal increase in water temperature of seasonally ice-covered lakes is projected to take place during the spring (Schindler et al., 1990; Winslow et al., 2017), concurrent with expected seasonal increases in air temperature during winter in the northern hemisphere (Christensen et al., 2007; Sharma et al., 2019). Such changes are likely to challenge aquatic organisms with life-history traits that include critical life stages during the winter-to-spring transition in these regions.

Warming spring water temperatures and increases in the length of the frost-free season can prolong annual growing seasons for aquatic organisms through warmer summers, longer and warmer autumns, and shorter ice-cover duration (Sharma et al., 2019, 2020). Temperature is an abiotic master factor for aquatic ecosystems because water temperature directly affects the physical and chemical properties of water, and phenology, reproductive events, metabolic rates, growth, and survival of aquatic organisms (Fry, 1964; Brett, 1979; Gillooly et al., 2002; Brown et al., 2004; Ohlberger et al., 2007; Busch et al., 2012; Cline et al., 2013; Little et al., 2020). The responses of many lake organisms to climate-derived changes in lake ecology are projected to be inadequate to counter the speed and magnitude of climate change, leaving some species vulnerable to decline and extirpation (Hoffmann & Sgrò, 2011). These pressures present challenges for biodiversity conservation and sustainability of ecosystem services. To navigate challenges, a foundational understanding of the primary threats to aquatic ecosystems and organisms across a range of spatial scales from local to global is needed (Vörösmarty et al., 2010; Halpern et al., 2015; Langhans et al., 2019).

The effects of increasing temperature on lake fishes are predicted to lead to declines in cold-water species and increases in warm-water species (Comte et al., 2013; Hansen et al., 2017). Species that possess narrow optimal thermal ranges, live near their thermal limits, or have long development times at cold temperatures are at-risk under warming climate scenarios as temperature can have strong direct and indirect effects at early-life stages (Blaxter, 1991; Pepin, 1991; Ficke et al., 2007; Mari et al., 2016; Lim et al., 2017; Dahlke et al., 2020). Unlike their marine counterparts, most freshwater fishes are restricted to their lake system, where their ability to evade the effects of climate change is impeded due to the isolated nature of lakes (Ficke et al., 2007). The ability of lake fish populations living close to their upper thermal limits to respond either through physiological adaptation or physiological plasticity will be required if species are to persist under increasingly stressful thermal conditions (Somero, 2009; Woolsey et al., 2015; Howells et al., 2016). Additionally, the amount of genetic variability within a population may constrain the ability to cope with environmentally-induced changes (*i.e.,* phenotypic plasticity; Somero, 2009; Schindler et al., 2015, 2010). Freshwater whitefishes, Salmonidae Coregoninae (hereafter coregonines), are of great socio-economic value (Nyberg et al., 2001; Ebener et al., 2008; Vonlanthen et al., 2009, 2012; Lynch et al., 2015, 2016), and are also considered to be critically sensitive to the effects of climate change because they are cold, stenothermic fishes (Stockwell et al., 2009; Elliott & Bell, 2011; Jeppesen et al., 2012; Isaak, 2014; Jonsson & Jonsson, 2014; Karjalainen et al., 2015, 2016a).

Coregonine fisheries worldwide have experienced population declines due to highly variable and weak year-class strengths (Nyberg et al., 2001; Vonlanthen et al., 2012; Anneville et al., 2015; Myers et al., 2015). In the 20th century, causes of decline included fishing and stocking practices (Anneville et al., 2015) and eutrophication causing poor incubation conditions (Müller, 1992; Vonlanthen et al., 2012). Today, in the northern hemisphere, the trophic state of lakes and fisheries management practices are improving, but coregonines continue to be the focus of reintroduction, restoration, and conservation efforts in many lakes (Favé & Turgeon, 2008; Zimmerman & Krueger, 2009; Bronte et al., 2017). Reasons for declining recruitment are unknown, but climate change, increasing water temperatures, and habitat degradation are hypothesized as the main causal factors (Nyberg et al., 2001; Marjomäki et al., 2004; Jeppesen et al., 2012; Anneville et al., 2015; Karjalainen et al., 2015, 2016a).

Coregonines generally spawn during late-autumn, embryos incubate over winter, and hatch in early- or late-spring (Stockwell et al., 2009; Karjalainen et al., 2015). The time between fertilization and hatching is inversely related to water temperature (Colby & Brooke, 1970, 1973; Luczynski & Kirklewska, 1984; Pauly & Pullin, 1988; Karjalainen et al., 2016a). Rising spring water temperatures (*e.g.,* > 4°C) trigger hatching of autumn-spawned coregonine embryos (Häkkinen et al., 2002; Urpanen et al., 2005; Karjalainen et al., 2015). The length of the newly-hatched larvae is negatively correlated with the temperature of incubation (Colby & Brooke, 1970, 1973; Luczynski & Kirklewska, 1984; Karjalainen et al., 2015). The long period between spawning and hatching exposes coregonines to a variety of thermal conditions, potentially resulting in a wide range of environmentally-induced phenotypes or plastic responses (Karjalainen et al., 2015, 2016a, 2016b). Coregonines are known to be behaviorally and developmentally plastic (Muir et al., 2013) and some species (*e.g.,* vendace *Coregonus albula* and European whitefish *C. lavaretus*) can respond to short- and long-winter conditions within the limits of phenotypic plasticity and through genetic adaptive changes, such as different embryo developmental rates (Karjalainen et al., 2015, 2016a).

Geographic variation is also important to consider with phenotypic plasticity. Many fishes in high-latitudes are adapted to relatively colder waters, extensive periods of ice cover, and strong seasonal daylight variations (Reist et al., 2006). Thus, high-latitude populations can show differential long-term adaptation to climates, compared to low-latitude population, across a latitudinal gradient (Conover & Present, 1990; Yamahira & Conover, 2002; Chavarie et al., 2010; Wilder et al., 2020). For example, a number of fishes have demonstrated an inverse relationship between length of the growing season and reproductive characteristics (*i.e.,* countergradient variation; Billerbeck et al., 2000; Chavarie et al., 2010; Conover & Present, 1990; Conover & Schultz, 1995; Jonassen, 2000; Schultz et al., 1996, 1998; Yamahira & Conover, 2002). Fishes at high latitudes experience lower temperatures overall and shorter growing seasons and should exhibit lower standard metabolic rates, growth rates, and smaller size-at-age than individuals at low latitudes (Reist et al., 2006; White et al., 2012). However, for cold-water stenothermic fishes, water temperatures at low latitudes may exceed their optimal range for significant portions of the growing season, or the amount of optimal thermal habitat decrease, while water temperatures at high latitudes may remain near the optimum for maximal growth efficiency throughout the growing season (Conover & Schultz, 1995). Because water temperature has a great influence on fish physiology and varies across latitudes, a wide range of adaptive responses by populations to increasing temperatures across latitudes is possible (Reist et al., 2006). However, in the absence of genetic information, phenotypic changes are difficult to distinguish as genetically based or the result of phenotypic plasticity (Merilä & Hendry, 2014; Fox et al., 2019). Large-scale experimental studies across geographic regions can aid our understanding of the role of phenotypic plasticity relative to adaptive evolution by leveraging long-term adaptations to distinct geographic locations and environments (Hoffmann & Sgrò, 2011; Merilä & Hendry, 2014; Fox et al., 2019). Coregonines occur broadly across northern latitudes and are an ideal group to test how cold-water lake fishes may respond to climate-driven shifts in environmental variables, such as water temperature.

Our objective was to compare the reaction norms of coregonine embryos within and among species from multiple sampling locations across North America and Europe to a standardized thermal gradient during incubation. We hypothesized that coregonines have differential levels of phenotypic plasticity in developmental and morphological traits of embryos in response to warming winter incubation conditions based on putative adaptation to their local environments. We predicted coregonines that share the same thermal environment respond similarly and geographically distinct groups with different thermal environments respond dissimilarly to increasing incubation temperatures.

## MATERIALS AND METHODS

### Study Species and Locations

We used a cross-lake, cross-continent, cross-species approach to evaluate the responses and thermal tolerances of coregonine embryos to changing thermal regimes. Wild-caught populations of cisco (*C. artedi*) in Lake Superior (LS-Cisco; USA/Canada) and Lake Ontario (LO-Cisco; USA/Canada), and vendace (LK-Vendace) and European whitefish (LK-Whitefish) in Lake Southern Konnevesi (Finland; Fig. 1) were sampled.

**Fig. 1.**
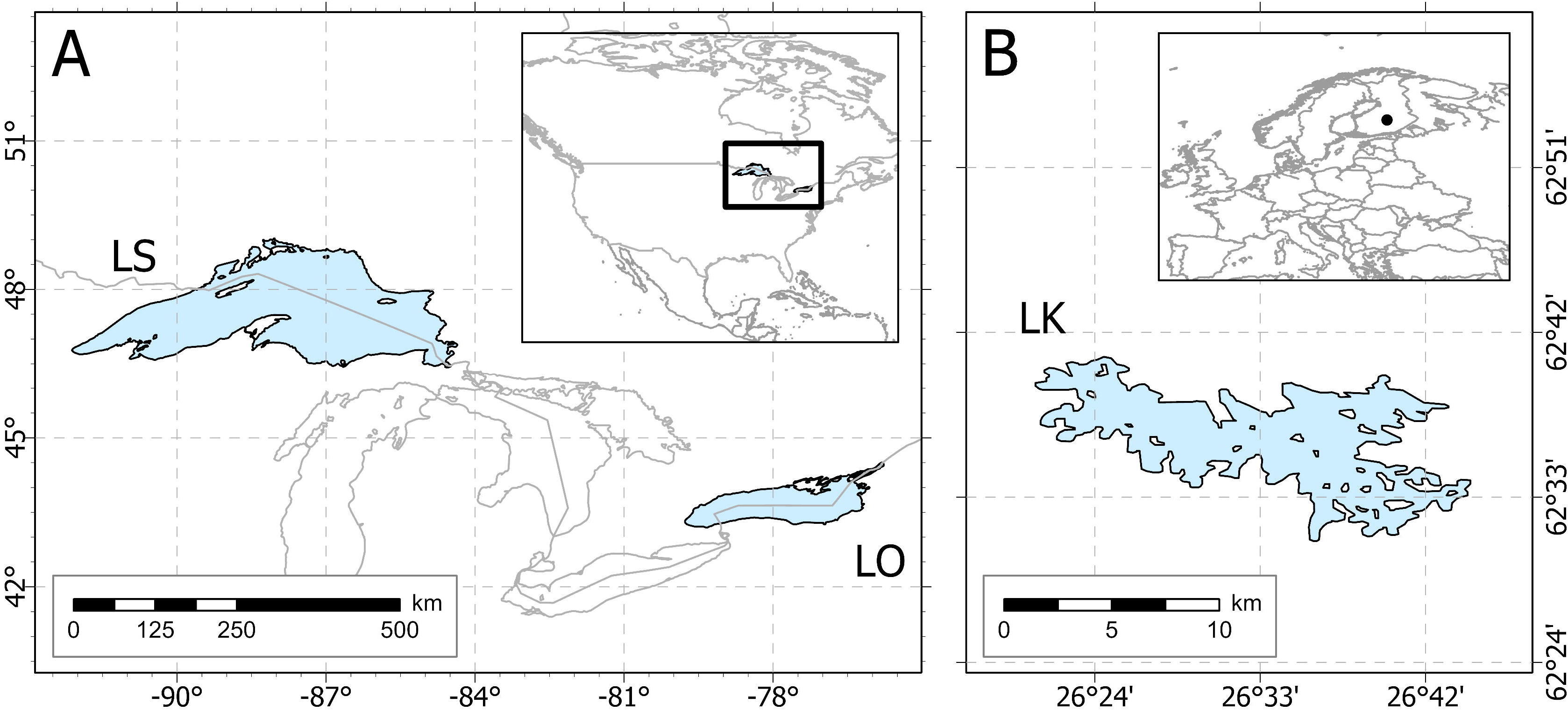
Map showing the location of each lake (LS = Lake Superior; LO = Lake Ontario; LK = Lake Southern Konnevesi) sampled in North America (A) and Europe (B)

Cisco is one of the most widespread of the North American species of coregonines (Eshenroder et al., 2016) and were one of the most abundant fish in the Great Lakes (Yule et al., 2013). Cisco is found in north-central to eastern United States and throughout most of Canada, with the lower Great Lakes close to its southernmost extent (Scott & Crossman, 1973). Cisco spawning is initiated when water temperatures decrease to 4-5°C in late Autumn (Pritchard, 1931; Eshenroder et al., 2016) and occurs at different spawning depths. Spawning can occur at depths ranging from 1-5 m in Lake Ontario (Pritchard, 1931; Paufve, 2019) and 10-64 m in Lake Superior (Dryer & Beil, 1964; Paufve, 2019). Experimental thermal optima for normal cisco embryo development is between 2 and 8°C (Colby & Brooke, 1970; Brooke & Colby, 1980). However, temperature data at historical spawning grounds indicate that *in-situ* incubations typically occur between 1 and 4°C, with Lake Ontario warmer than Lake Superior (Fig. 2; unpublished data). Optimal growth and upper lethal temperatures for cisco adults are estimated to be 18° and 26°C, respectively (Edsall & Colby, 1970; McCormick et al., 1971; Jobling, 1981).

**Fig. 2.**
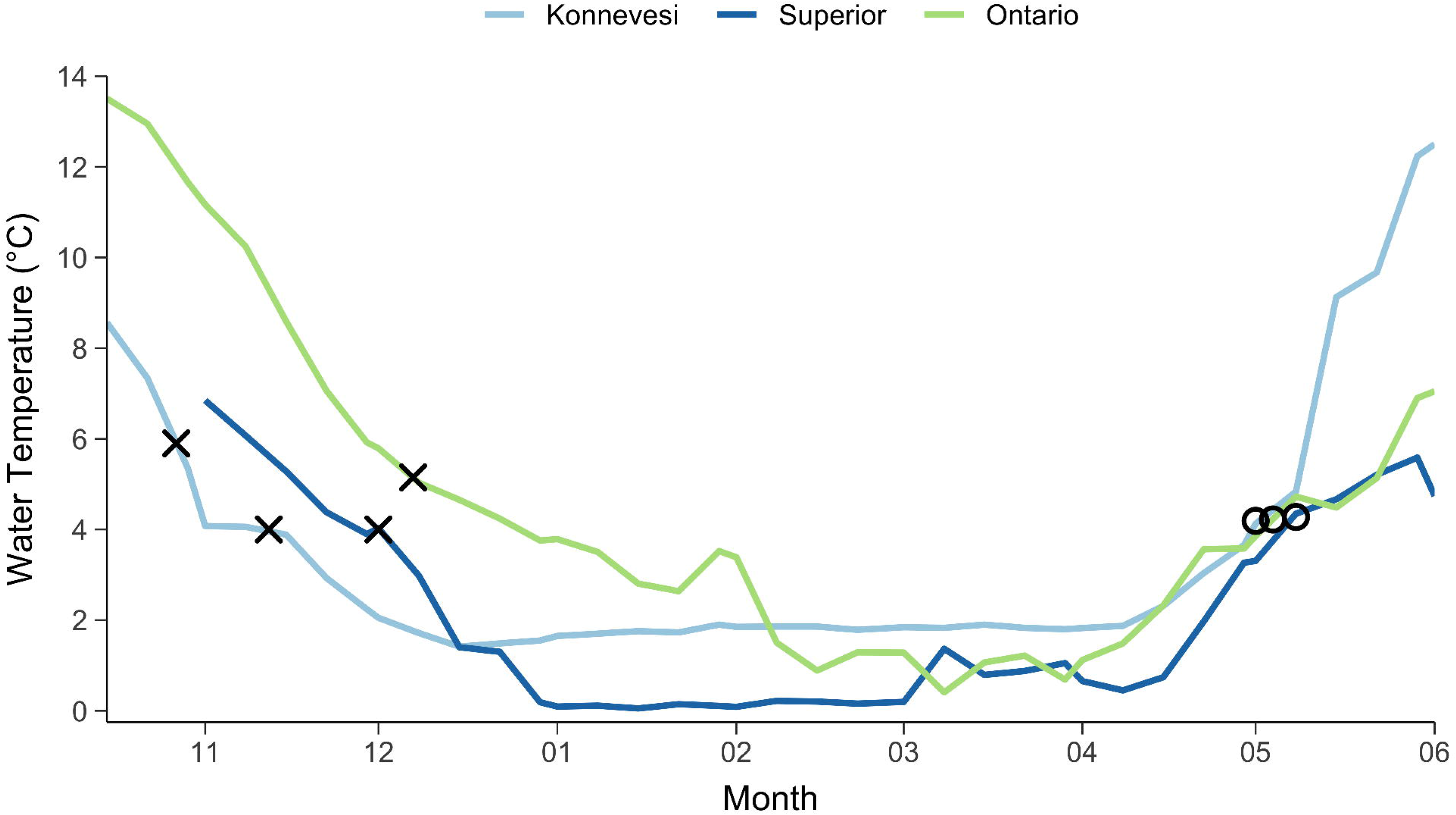
Winter water temperatures from Lake Southern Konnevesi, Lake Superior, and Lake Ontario. Lake Southern Konnevesi and Lake Superior data were recorded using *in-situ* sensors on the lakebed (10-m deep). Lake Ontario data was recorded using remote sensing sea surface temperatures. X indicates *ca.* spawning and O indicates *ca.* hatching. The earlier spawning (X) in Lake Southern Konnevesi indicates vendace (*Coregonus albula*) and latter for European whitefish (*C. lavaretus*)

Vendace and European whitefish are widely distributed in Northern Europe (Sipponen et al., 2006). Vendace spawning is initiated when water temperatures decrease to 6°C at the end of October in Lake Southern Konnevesi and lasts 2-3 weeks (Karjalainen & Marjomäki, 2018). European whitefish spawn 2-3 weeks later in November when water temperatures decrease to 4- 5°C (Karjalainen et al., 2015). Spawning of vendace occurs in the littoral and sublittoral zone of lakes and eggs dispersed widely at depths less than 20 m (Heikkilä et al., 2006; Karjalainen et al., In press). Whitefish are known to spawn at shallower depths than vendace. Embryos are incubated at 1-2°C until the beginning of April, when water temperatures have gradually increased, and hatching peaks after ice-out at 4-6°C (Fig. 2, Karjalainen et al., 2015). Although whitefish spawn later than vendace, their hatching occurs at the same time or slightly earlier than vendace (Karjalainen et al., 2015). Optimal growth and upper lethal lethal temperatures for adults are estimated to be 18° and 27°C for immature vendace and 18° and 29°C for European whitefish, respectively (Tapaninen et al., 1998, Vielma et al. 2002, Siikavoipio et al. 2013).

### Adult Collections

Adults were sampled using multi-mesh gillnets (51-89 mm stretched mesh) in Lake Superior (46.85°, -90.55°), trap nets in Lake Ontario (44.05°, -76.20°), and seines (vendace) and multi-mesh gillnets (European whitefish; 27-35 mm knot to knot mesh) in Lake Southern Konnevesi (62.58°, 26.58°). Adult field collections occurred during coregonine spawning periods for Lake Ontario and Lake Superior. On Lake Southern Konnevesi, adults were collected prior to spawning and held in aquaculture pools with water fed directly from the lake until spawning was initiated. Demographic data (*e.g.,* total length, mass, and egg diameter) were collected on sampled adults. All sampling, fertilization, and experimental work for study groups on each continent were conducted at a single laboratory in North America (University of Vermont (UVM), USA) and Europe (University of Jyväskylä (JYU), Finland). Experiments were performed during the 2018-19 season in Finland and the 2019-20 season in the USA.

For clarity, our operational use of a study group is to represent a single species haphazardly sampled from a single location within a single lake (*e.g.,* cisco from the Apostle Islands region in Lake Superior). Our sampling efforts represent a single location within large lakes and likely do not capture all possible genetic variation within a species or population.

### Crossing Design and Fertilization

Eggs and milt were stripped from 12 females and 16 males from each study group and artificially fertilized under a blocked, nested full-sib, half-sib fertilization design to create a maximum of 48 full-sibling families nested within half-siblings per group. The crossing design maximized the amount of genetic variation and minimized the potential loss of multiple families if a female or male produced poor quality gametes, for a given total number of families, compared to a full-factorial design. Adults used in the experiment were divided into three or four fertilization blocks. A single block consisted of four males each paired to three females, where all offspring of a given female were full siblings. Fertilizations were performed block by block to ensure germ cell survival.

Approximately 200 eggs per female were fertilized with an equal amount of milt (5-15 μl) from each male in the block. After the addition of milt, water was added to activate the germ cells and gently mixed for one minute. The embryos were rinsed with water 2-3 times until the water was clear. Temperature of the water used during all fertilizations was ca. 4°C. Reconstituted freshwater medium (ISO 6341, 2012) was used during fertilizations to standardize the chemical properties of the water used among study groups and between labs. Embryos were transported in coolers either by shipping overnight for Lake Superior or driven same-day for Lake Ontario. A temperature logger (HOBO^®^ Water Temperature Pro v2) recorded air temperature inside the cooler during transport (Lake Superior: mean (SD) = 2.80°C (0.21); Lake Ontario: mean (SD) = 3.28°C (0.37)). No embryo transport was required for Lake Southern Konnevesi. Fertilization success was determined by haphazardly taking 10 embryos from each family and assessing under microscopy within 72-hours post-fertilization (Oberlercher & Wanzenböck, 2016). If fertilization was low (<30%), the family was removed from the experimental setup.

### Rearing Conditions

Embryos from successfully fertilized families were individually distributed into 24-well cell culture microplates and incubated in 2 ml of reconstituted freshwater. Reconstituted freshwater was used during incubation to maintain sterility, prevent bacterial growth in the wells, and eliminate the need for fungicide treatments on the embryos. A total of 36 embryos per family were used for Lake Southern Konnevesi species and 48 embryos per family for each of Lake Ontario and Lake Superior cisco. Families were randomly distributed across three or four microplates (*i.e.,* 12 eggs per family per microplate and two families per 24-well microplate). Microplates from each study group were incubated at target constant temperatures of 2.0 (coldest), 4.5 (cold), 7.0 (warm), and 9.0°C (warmest) and randomly placed in climate-controlled chambers at UVM (Memmert^®^ IPP260Plus) and climate-controlled rooms at JYU (Huurre^®^).

The range of experimental incubation temperature treatments was chosen to mimic *in-situ* mean temperatures and to exceed optimum embryonic development temperatures. Forced airflow was used in both the climate-controlled chambers and rooms to ensure equal air circulation around the microplates. All microplates were covered to minimize evaporation. Microplate orientation and position were rotated weekly to eliminate any temperature heterogeneity within the chambers and rooms. HOBO^®^ TMCx-HD temperature probes were used at UVM to record water temperatures hourly in 50-ml beakers placed inside each climate chamber, while Escort iMini temperature loggers were used directly inside the well of a subset of microplates for each temperature treatment at JYU. Daily mean water temperatures were calculated. Incubations took place in the dark, with the exception of short (< 60 mins) maintenance periods. Microplates were checked weekly for dead eggs and the eye-up stage. During the hatch period, microplates were checked on a two-day cycle for newly hatched larvae. For cisco, all newly hatched larvae were photographed for developmental and morphological traits (Nikon^®^ D5600 and Nikon^®^ AF-S DX 18-55mm lens). Egg size, total length, and yolk-sac axes were measured from images using Olympus^®^ LCmicro. For Lake Southern Konnevesi, the larvae were preserved in ethanol at hatch and flushed and soak in distilled water for 15 min before measuring the total length and fresh mass under the microscope (Karjalainen, 1992).

Mean water temperature during incubations was maintained near the target incubation temperature for the cold and warm treatments at each lab. Mean incubation water temperatures for the cold and warmest treatments were lower than the target incubation temperature at JYU, but not at UVM (Table 1).

**Table 1.** Mean (SD) water temperatures during embryo incubations at the University of Vermont (UVM) and University of Jyväskylä (JYU).

| Laboratory | Incubation Temperature Treatment (°C) |  |  |  |
| --- | --- | --- | --- | --- |
|  | 2.0<br>Coldest | 4.5<br>Cold | 7.0<br>Warm | 9.0<br>Warmest |
| UVM | 2.0 (0.5) | 4.4 (0.2) | 6.9 (0.2) | 8.9 (0.3) |
| JYU | 2.2 (1.5) | 4.0 (0.7) | 6.9 (0.5) | 8.0 (0.6) |

### Developmental and Morphological Traits

Embryo survival was estimated as the percent of embryos surviving between the eye-up and hatch stages to rule out unfertilized eggs which can bias survival estimates. Incubation period was assessed by two variables: the number of days from fertilization to hatching (days post-fertilization; DPF) and the sum of the degree-days (accumulated degree-days; ADD). Total length-at-hatch (mm) and yolk-sac volume (YSV; mm^3^) were measured from five individuals per family at, or as close as possible to, 50% hatching for each family. Yolk-sac volume was calculated assuming the shape of an ellipse (Blaxter & Hempel, 1963):

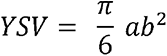

where a = length of the yolk sac (mm) and b = hei^6^ght of the yolk sac (mm).

### Statistical Analyses and Estimation of Variance Components

Embryo survival was analyzed as a binomial response variable, and incubation period, length-at-hatch, and yolk-sac volume at hatching as continuous response variables. Early embryo mortality and variable fertilization success induced from fertilization failure produced inequalities in the number of offspring among families. The sample size for incubation period is a function of embryo survival and subsequently resulted in an unbalanced design. All non-proportional data were checked visually for approximate normality using histograms and Q-Q plots before the analysis with parametric models. A cubic transformation was applied to LAH for cisco and a cubic root transformation was applied to DPF, ADD, and YSV to normalize the distributions.

Therefore, binary data *(i.e.,* embryo survival) were analyzed with binomial generalized linear mixed-effects models (LMM) and transformed variables (*i.e.,* DPF, ADD, LAH, and YSV) were analyzed with restricted maximum likelihood LMMs with the *lme4* package v.1.1-26 (Bates et al., 2015). To eliminate any confounding effects between continents, cisco from lakes Superior and Ontario were analyzed independently from vendace and European Whitefish from Lake Southern Konnevesi. Population (for cisco only), species (for Lake Southern Konnevesi only), and incubation temperature were included as fixed effects and female, male, family (female and male combination), microplate, and fertilization block as random effects. Because embryos were raised independently, the replication unit in the statistical models is the individual embryo.

Although incubation temperature was treated as a fixed variable based on our experimental design (*i.e.,* few treatment levels with high replication), we acknowledge that temperature is a continuous, independent variable in nature. All traits were examined for population or species, depending on the continent, and incubation temperature effects in addition to individual parental effects (female, male, and family effects), microplate, fertilization block, and all possible interactions with backward, stepwise effect-selection using the *buildmer* package v.1.7.1 (Voeten, 2020). The maximal model for each trait was selected by comparing a model including or lacking the term of interest to the reference model based on changes in log-likelihood, Akaike information criterion, Bayesian information criterion, and change in explained deviance.

Significance values for the mixed-effects model parameters (*i.e.,* population, species, incubation temperature, interaction effects, and any random-effects selected) were determined using a likelihood ratio test between the maximal model and reduced models with the model parameter of interest removed (Myers et al. 1995; Luke 2017). All statistical tests used L = 0.05 to determine significance.

To allow for interspecific comparisons, the response to temperature for each trait was standardized to the optimal temperature for each study group. Based on literature data (cf. above), the coldest incubation temperature treatment (2.0°C and 2.2°C; Table 1) was assumed to be the optimal incubation temperature. For each trait, the within-family mean was calculated for each temperature treatment and the percent change from the optimal temperature estimated. Standard error was calculated as the among-family variation in percent change.

The phenotypic variance components were partitioned into random effects for female, male, female x male, and unexplained or random residual variance components using mixed-effects models with the *fullfact* package v.1.3 (Houde & Pitcher, 2016) for each study group and incubation temperature treatment. Negative variance components were treated as zero (Neff & Pitcher, 2005). The percent of total phenotypic variation was used to calculate the Pearson correlation coefficient between each variance component and the increase in incubation temperature for each study group. A threshold of ±0.7 (*i.e.,* an R^2^ of 0.49) was used to categorize correlations either positive or negative, with all values in between as no correlation. European whitefish from Lake Southern Konnevesi were removed from this analysis due to a low embryo survival and a low number of families. All analyses were performed in R version 4.0.3 (R Core Team, 2020).

## RESULTS

### Spawning Adults

Total lengths and fresh mass of spawning adults used for gamete collection varied widely among study groups (Table 2). LK-Vendace were notably smaller than all other study groups. The remaining study groups varied less in size, but LK-Whitefish were smaller than LS-Cisco and LO-Cisco.

**Table 2.**
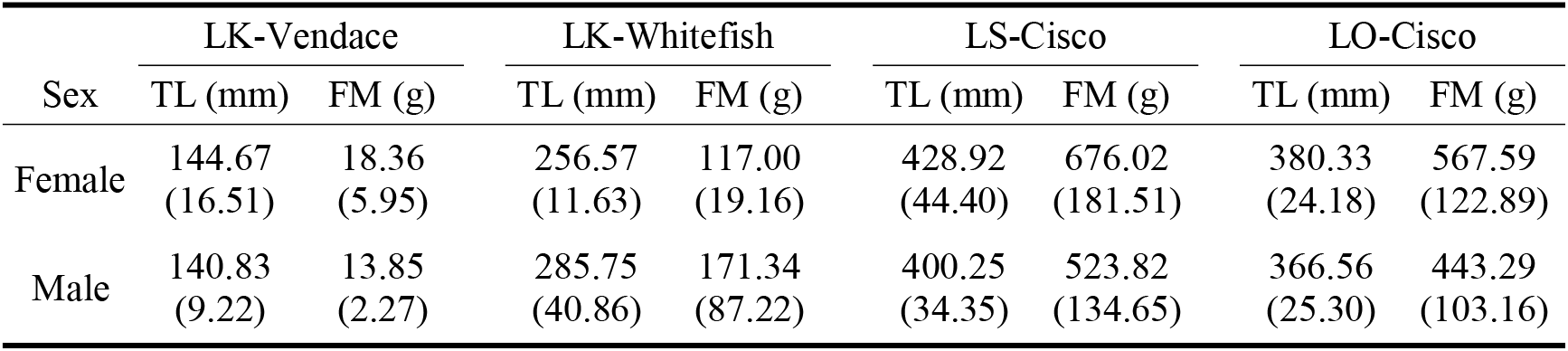
Mean (SD) total length (TL) and fresh mass (FM) of the female and males from Lake Southern Konnevesi (LK-Vendace (*Coregonus albula*) and LK-Whitefish (*C. lavaretus*)), Lake Superior (LS-Cisco (*C. artedi*)), and Lake Ontario (LO-Cisco).

| Sex | LK-Vendace |  | LK-Whitefish |  | LS-Cisco |  | LO-Cisco |  |
| --- | --- | --- | --- | --- | --- | --- | --- | --- |
|  | TL (mm) | FM (g) | TL (mm) | FM (g) | TL (mm) | FM (g) | TL (mm) | FM (g) |
| Female | 144.67<br>(16.51) | 18.36<br>(5.95) | 256.57<br>(11.63) | 117.00<br>(19.16) | 428.92<br>(44.40) | 676.02<br>(181.51) | 380.33<br>(24.18) | 567.59<br>(122.89) |
| Male | 140.83<br>(9.22) | 13.85<br>(2.27) | 285.75<br>(40.86) | 171.34<br>(87.22) | 400.25<br>(34.35) | 523.82<br>(134.65) | 366.56<br>(25.30) | 443.29<br>(103.16) |

The LK-Vendace females had the smallest egg diameters and LO-Cisco females had the largest egg diameters among the study groups (Table 3). LK-Whitefish and LS-Cisco egg diameters were similar (Table 3).

**Table 3.** Mean (SD) egg diameter of females with the number of eggs measured (N) from Lake Southern Konnevesi (LK-Vendace (*Coregonus albula*) and LK-Whitefish (*C. lavaretus*)), Lake Superior (LS-Cisco (*C. artedi*)), and Lake Ontario (LO-Cisco).

| Population | Egg Diameter (mm) | N |
| --- | --- | --- |
| LK-Vendace | 1.58 (0.11) | 273 |
| LK-Whitefish | 2.13 (0.12) | 70 |
| LS-Cisco | 2.14 (0.12) | 140 |
| LO-Cisco | 2.30 (0.08) | 240 |

### Developmental and Morphological Traits and Variance Components

All cisco traits had significant interaction effects between population and incubation temperature (maximum *P* < 0.001; Tables 4 and 5). All vendace and European whitefish traits had significant interaction effects between species and incubation temperature (maximum *P* = 0.002; Tables 4 and 5). The interaction effects precluded any interpretation of main effects, but did suggest different norms of reaction for the model groups. Below we describe the interaction effects. All random effects (*i.e.,* female, male, and female x male) were significant (maximum *P* = 0.038) except male for LAH and female x male for LAH and YSV in cisco (Tables 4 and 5). All statistical model results can be found in Tables 4 and 5.

**Table 4.**
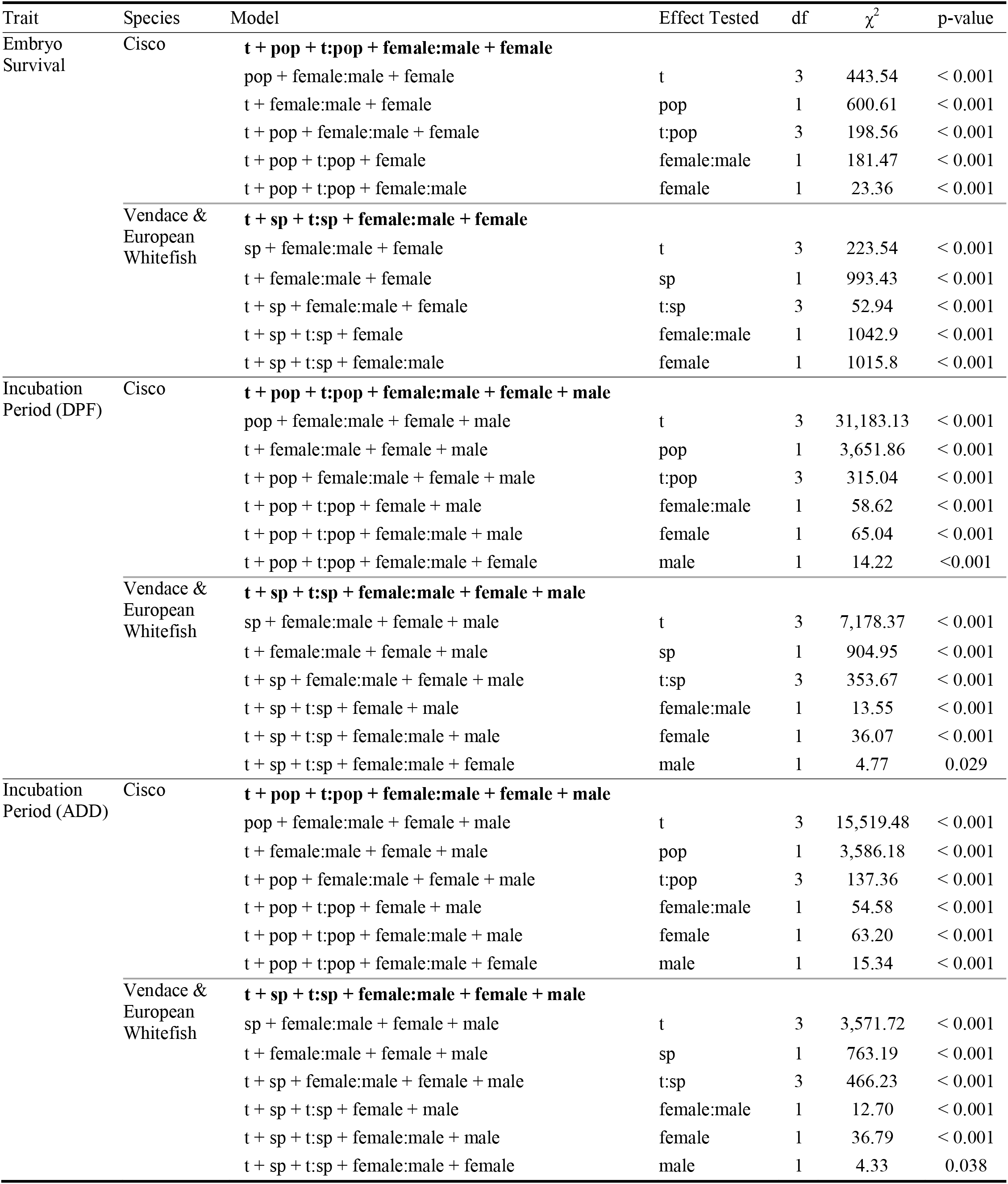
Likelihood ratio test output for each model selected for embryo survival and incubation period (number of days post-fertilization (DPF) and accumulated degree days (°C; ADD)) from lakes Superior and Ontario cisco (*Coregonus artedi*) and Lake Southern Konnevesi vendace (*C. albula*) and European whitefish (*C. lavaretus*). t indicates temperature, pop indicates population, and sp indicates species. The full model that was selected is bolded for each trait and species.

**Table 5.**
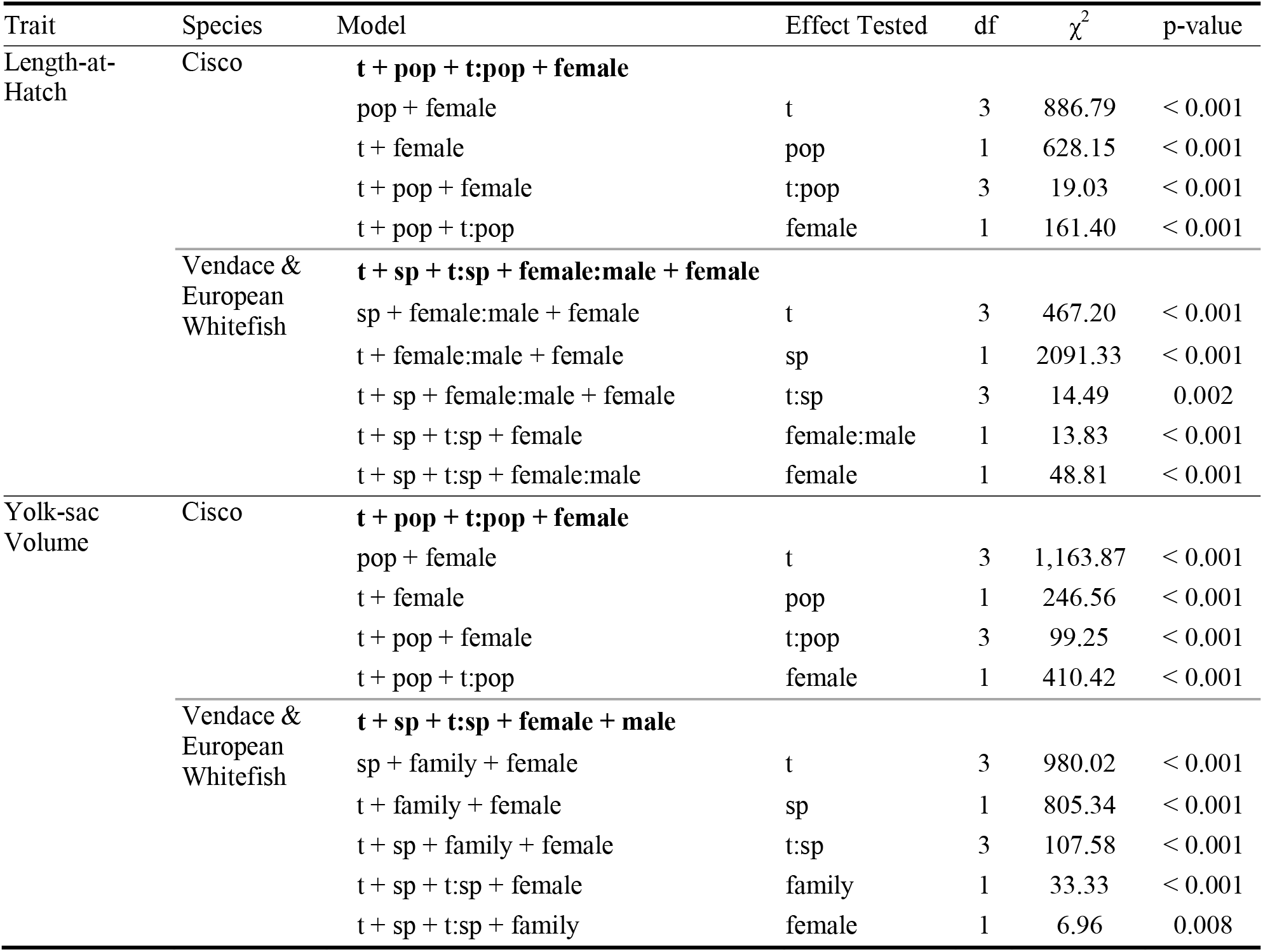
Likelihood ratio test output for each model selected for length-at-hatch (mm) and yolk-sac volume (mm^3^) from lakes Superior and Ontario cisco (*Coregonus artedi*), Lake Southern Konnevesi vendace (*C. albula*), and Lake Southern Konnevesi European whitefish (*C. lavaretus*). t indicates temperature and pop indicates population. The full model that was selected is bolded for each trait and species.

| Trait | Species | Model | Effect Tested | df | $\chi^2$ | p-value |
| --- | --- | --- | --- | --- | --- | --- |
| Length-at-Hatch | Cisco | **t + pop + t:pop + female** |  |  |  |  |
|  |  | pop + female | t | 3 | 886.79 | < 0.001 |
|  |  | t + female | pop | 1 | 628.15 | < 0.001 |
|  |  | t + pop + female | t:pop | 3 | 19.03 | < 0.001 |
|  |  | t + pop + t:pop | female | 1 | 161.40 | < 0.001 |
|  | Vendace & European Whitefish | **t + sp + t:sp + female:male + female** |  |  |  |  |
|  |  | sp + female:male + female | t | 3 | 467.20 | < 0.001 |
|  |  | t + female:male + female | sp | 1 | 2091.33 | < 0.001 |
|  |  | t + sp + female:male + female | t:sp | 3 | 14.49 | 0.002 |
|  |  | t + sp + t:sp + female | female:male | 1 | 13.83 | < 0.001 |
|  |  | t + sp + t:sp + female:male | female | 1 | 48.81 | < 0.001 |
| Yolk-sac Volume | Cisco | **t + pop + t:pop + female** |  |  |  |  |
|  |  | pop + female | t | 3 | 1,163.87 | < 0.001 |
|  |  | t + female | pop | 1 | 246.56 | < 0.001 |
|  |  | t + pop + female | t:pop | 3 | 99.25 | < 0.001 |
|  |  | t + pop + t:pop | female | 1 | 410.42 | < 0.001 |
|  | Vendace & European Whitefish | **t + sp + t:sp + female + male** |  |  |  |  |
|  |  | sp + family + female | t | 3 | 980.02 | < 0.001 |
|  |  | t + family + female | sp | 1 | 805.34 | < 0.001 |
|  |  | t + sp + family + female | t:sp | 3 | 107.58 | < 0.001 |
|  |  | t + sp + t:sp + female | family | 1 | 33.33 | < 0.001 |
|  |  | t + sp + t:sp + family | female | 1 | 6.96 | 0.008 |

#### Embryo Survival

Embryo survival was highest among all study groups at the coldest temperature and lowest at the warmest temperature (Fig. 3). The effect of temperature for cisco depended on population because embryo survival was higher for LO-Cisco (99.3%) than LS-Cisco (80.0%) at the coldest temperature but not different between populations (difference < 0.1%) at the warmest temperature (Fig. 3). For Lake Southern Konnevesi, the effect of temperature depended on species because the difference in embryo survival between LK-Vendace and LK-Whitefish was less pronounced at the coldest temperature (29.0%) than at the warmest temperature (50.5%; Fig. 3). LK-Vendace and LK-Whitefish embryo survival had a differential temperature response as LK-Whitefish had a greater decrease (74.4%) than LK-Vendace (17.7%) from the coldest to warmest incubation temperatures. LK-Whitefish had the strongest, decreasing response to increasing incubation temperatures compared to all other study groups (Fig. 3).

**Fig. 3.**
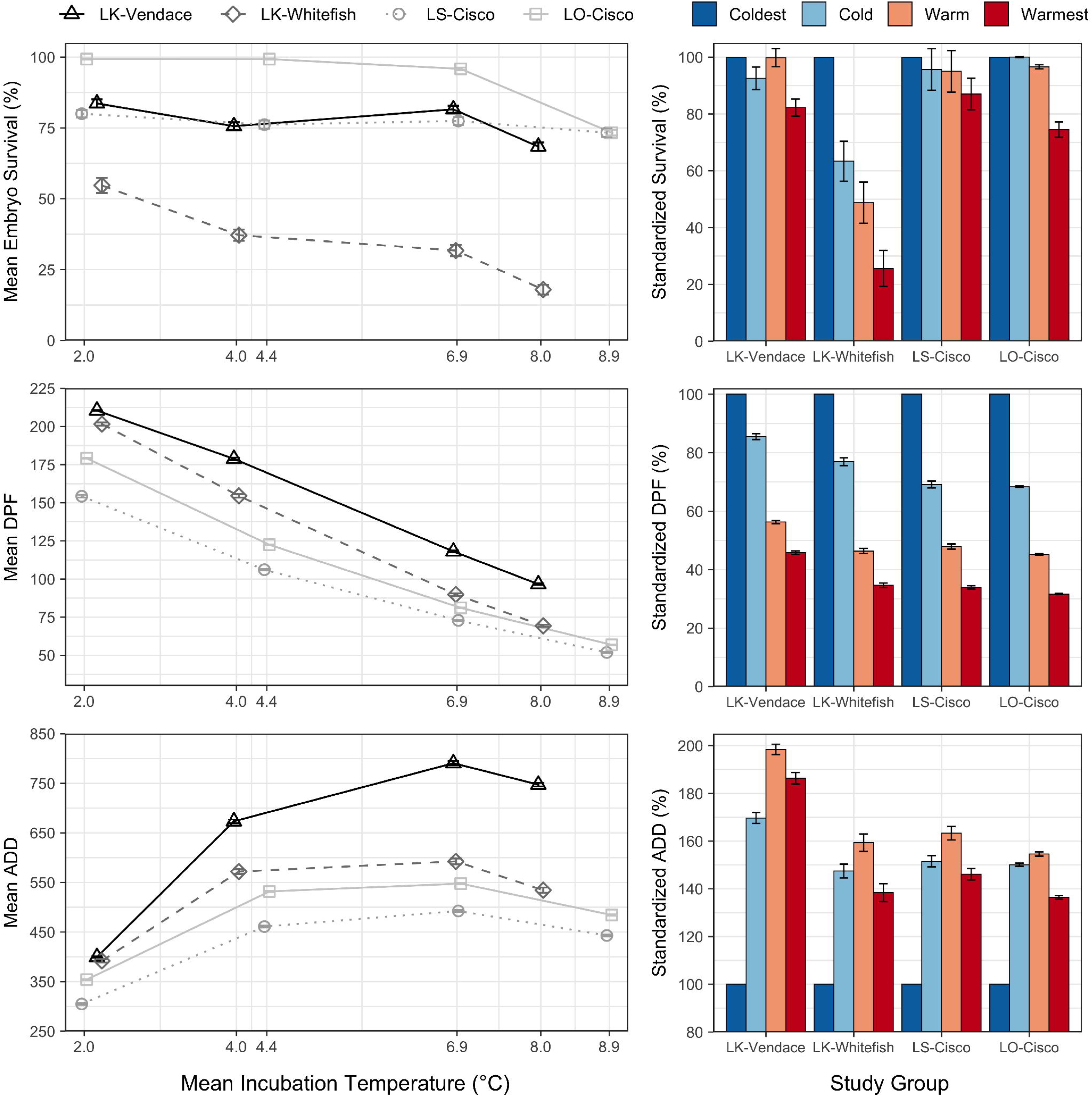
Mean embryo survival (%) and incubation period (number of days post-fertilization (DPF) and accumulated degree days (°C; ADD)) at each incubation temperature (°C; left) and standardized temperature responses within each study group (%; right) from Lake Southern Konnevesi (LK-Vendace (*Coregonus albula*) and LK-Whitefish (*C. lavaretus*)), Lake Superior (LS-Cisco (*C. artedi*)), and Lake Ontario (LO-Cisco). Error bars indicate standard error

In the phenotypic variance component analysis, the residual error was the largest component of phenotypic variation in embryo survival (means > 55.2%) for all study groups (Fig. 4, Online Resource 1). The mean female variance had the highest percentage, excluding error, of the phenotypic variation in embryo survival for LK-Vendace (17.4%), LS-Cisco (24.1%), and LO- Cisco (19.9%; Fig. 4, Online Resource 1). The female variance component correlations for embryo survival had either negative or no correlations to increasing temperature; however, male and error variances had positive and no correlations suggesting that as the female component decreases at higher temperatures the importance of the male component and error increases (Table 6).

**Fig. 4.**
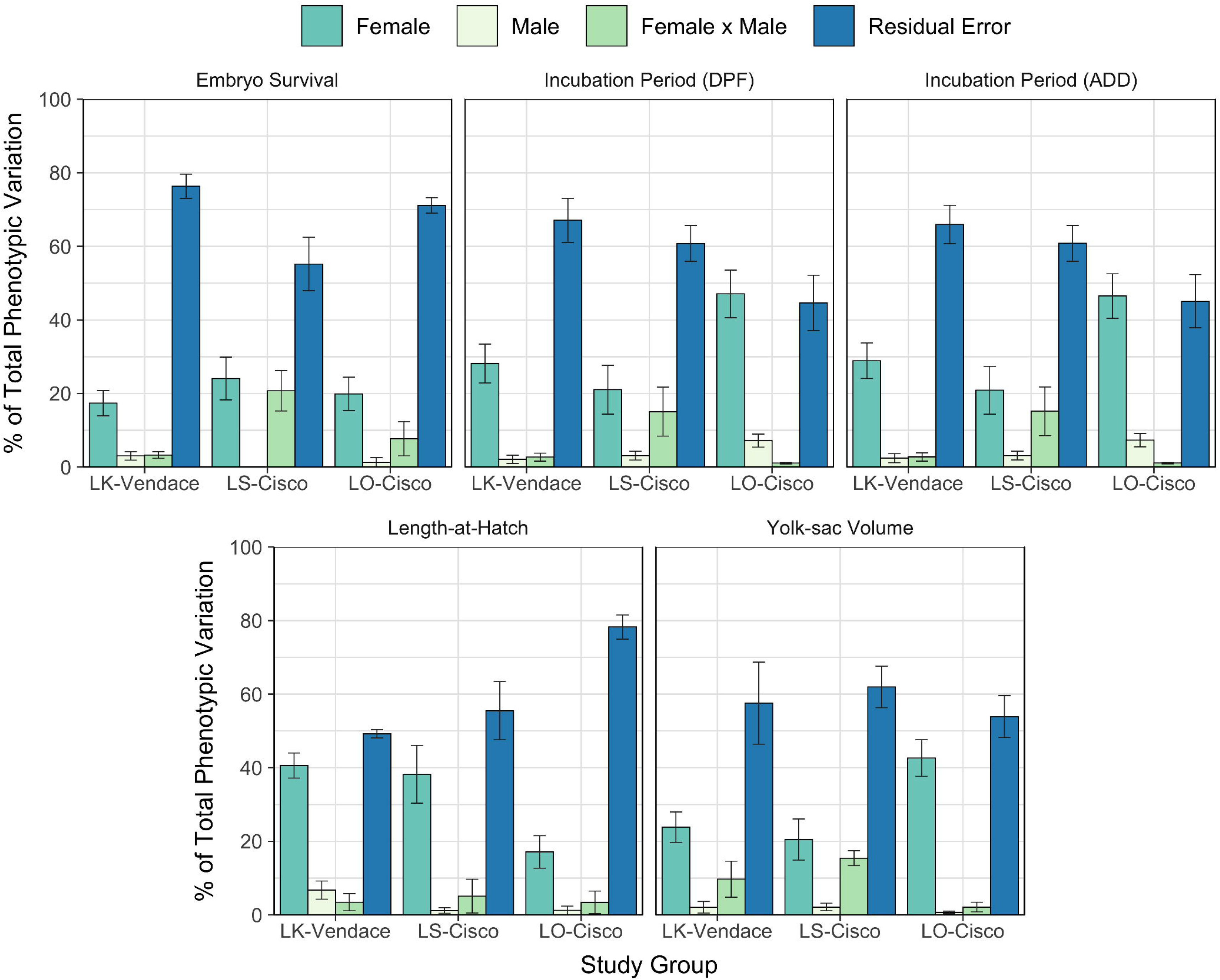
Mean percent of total phenotypic variation across incubation temperatures for embryo survival, incubation period (number of days post-fertilization (DPF) and accumulated degree days (°C; ADD)), length-at- hatch (mm), and yolk-sac volume (mm^3^) from Lake Southern Konnevesi vendace (LK-Vendace (*Coregonus albula*), Lake Superior cisco (LS-Cisco (*C. artedi*)), and Lake Ontario cisco (LO-Cisco). Error bars indicate standard error

**Table 6.** Phenotypic variation component correlation directions from increasing incubation temperature for embryo survival (%), incubation period (number of days post-fertilization; DPF), incubation period (accumulated degree-days; ADD), length-at-hatch (mm), and yolk-sac volume (mm^3^) from Lake Southern Konnevesi vendace (LK-Vendace (*Coregonus albula*)), Lake Superior cisco (LS-Cisco (*C. artedi*)), and Lake Ontario cisco (LO-Cisco). - indicates a negative correlation, + indicates a positive correlation, and 0 indicates no correlation.

| Trait | Study Group | Correlation Direction |  |  |  |
| --- | --- | --- | --- | --- | --- |
|  |  | Dam | Sire | Dam:Sire | Error |
| Embryo Survival | LK-Vendace | - | + | 0 | + |
|  | LS-Cisco | - | 0 | 0 | 0 |
|  | LO-Cisco | 0 | + | 0 | + |
| Incubation Period (DPF) | LK-Vendace | 0 | 0 | + | 0 |
|  | LS-Cisco | - | 0 | 0 | 0 |
|  | LO-Cisco | - | 0 | 0 | + |
| Incubation Period (ADD) | LK-Vendace | 0 | - | + | 0 |
|  | LS-Cisco | - | 0 | 0 | 0 |
|  | LO-Cisco | - | - | 0 | + |
| Length-at-Hatch | LK-Vendace | 0 | - | + | 0 |
|  | LS-Cisco | - | 0 | 0 | 0 |
|  | LO-Cisco | - | 0 | + | 0 |
| Yolk-sac Volume | LK-Vendace | + | 0 | + | - |
|  | LS-Cisco | - | + | - | + |
|  | LO-Cisco | 0 | 0 | 0 | 0 |

#### Incubation Period (days post-fertilization)

The number of days post-fertilization to hatching was highest for all study groups at the coldest temperature and decreased as temperature increased (Fig. 3). For cisco, DPF was higher for LO- Cisco (179.2 days) than LS-Cisco (154.3 days) at the coldest temperature and the difference between populations was less pronounced at the warmest temperature (difference = 5.0 days; Fig. 3). For Lake Southern Konnevesi, the effect of temperature depended on species because the difference in DPF between LK-Vendace and LK-Whitefish was less pronounced at the coldest temperature (8.9 days) than at warmest temperature (27.3 days; Fig. 3). All study groups had similar responses to temperature, with between 54.2 to 68.3% decreases in DPF from the coldest to warmest treatments. However, LS-Cisco, LO-Cisco, and LK-Whitefish had a greater decrease in DPF (66.1, 68.3, 65.3%, respectively), than LK-Vendace (54.2%; Fig. 3).

In the phenotypic variance component analysis, the residual error was the largest component of phenotypic variation in DPF (means >60.8%) for LK-Vendace and LS-Cisco (Fig. 4, Online Resource 1). The mean female variance was the largest phenotypic variation component in DPF for LO-Cisco (47.1%). LK-Vendace and LS-Cisco had similar mean female variances for DPF across all temperatures, with 28.1 and 21.0%, respectively (Fig. 4, Online Resource 1). The DPF correlations for female effect had a negative response to temperature for LS-Cisco and LO-Cisco (Table 6).

#### Incubation Period (accumulated degree-days)

Accumulated degree-days were highest for all study groups at 6.9°C (Fig. 3). The effect of temperature for cisco depended on population because ADD was higher for LO-Cisco (531.9 and 547.7 ADD) than LS-Cisco (461.0 and 492.5 ADD) at the cold and warm temperatures, respectively, and the differences between populations were less pronounced at the coldest and warmest temperatures (differences = 49.2 and 41.3 ADD, respectively; Fig. 3). LS-Cisco and LO-Cisco ADD responded similarly to increasing incubation temperature. For Lake Southern Konnevesi, the effect of temperature depended on species because the difference in ADD between LK-Vendace and LK-Whitefish was less pronounced at the coldest temperature (7.7 ADD) than at the warm temperature (198.1 ADD; Fig. 3). LK-Vendace and LK-Whitefish ADD had a differential temperature response as LK-Vendace had a greater increase (198.4%) than LK- Whitefish (159.4%) from the coldest to warm treatment. LK-Vendace had the strongest, increasing response to increasing incubation temperatures compared to all other study groups (Fig. 3).

In the phenotypic variance component analysis and correlations, ADD had a similar response as DPF among all study groups as the data only had a different temperature scaling factor (Fig. 4, Online Resource 1, Table 6).

#### Length-at-Hatch

All study groups had a common, decreasing response in LAH as temperature increased (Fig. 5). For cisco, LAH was higher for LO-Cisco (11.32 and 9.75 mm) than LS-Cisco (10.21 and 8.68 mm) at the coldest and warmest temperatures, respectively, and the difference between populations was less pronounced at the cold and warm temperatures (differences = 0.99 and 0.90 mm, respectively; Fig. 5). LS-Cisco and LO-Cisco responded to increasing incubation temperature with a 15.9 and 13.8% respective decrease in LAH from the coldest to warmest treatments. For Lake Southern Konnevesi, the effect of temperature depended on species because the difference in LAH between LK-Vendace and LK-Whitefish was more pronounced at the cold and warm temperatures (2.73 and 2.72 mm, respectively) than at the coldest and warmest temperatures (2.68 and 2.61 mm, respectively; Fig. 5). LK-Vendace and LK-Whitefish each responded similarly to temperature with a 9.0 and 9.2% respective decrease in LAH from the coldest to warmest treatments. LS-Cisco and LO-Cisco LAH had a stronger, decreasing response to increasing incubation temperatures than LK-Vendace and LK-Whitefish (Fig. 5).

**Fig. 5.**
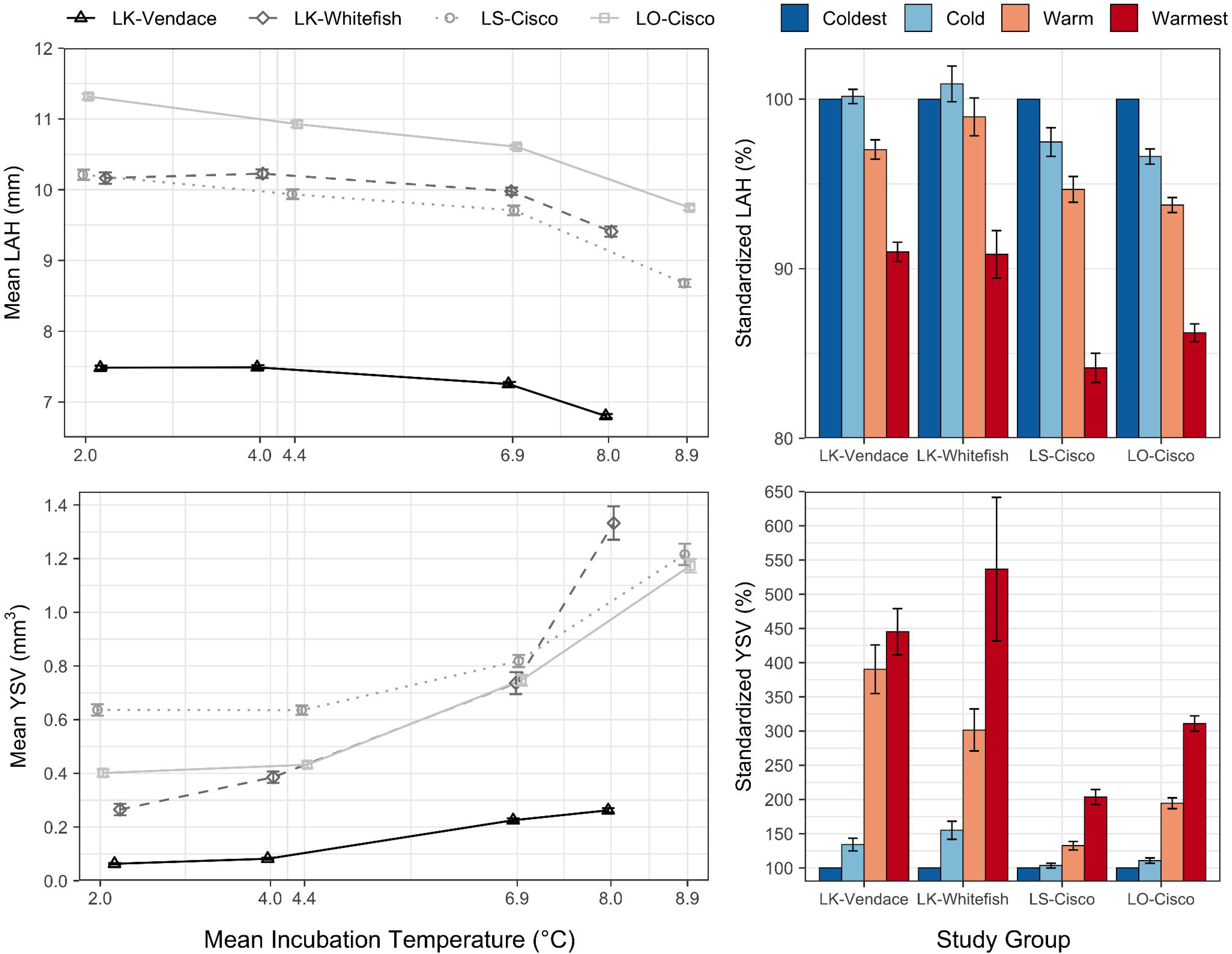
Mean length-at-hatch (mm; LAH) and yolk-sac volume (mm^3^; YSV) at each incubation temperature (°C; left) and standardized temperature responses within each study group (%; right) from Lake Southern Konnevesi (LK-Vendace (*Coregonus albula*) and LK-Whitefish (*C. lavaretus*)), Lake Superior (LS-Cisco (*C. artedi*)), and Lake Ontario (LO-Cisco). Error bars indicate standard error

In the phenotypic variance component analysis, the residual error was the largest component of phenotypic variation in LAH (means >49.2%) for all study groups (Fig. 4, Online Resource 2). The mean female variance had the highest percentage, excluding error, of the phenotypic variation in LAH for LK-Vendace (40.6%), LS-Cisco (38.2%), and LO-Cisco (17.1%; Fig. 4, Online Resource 2). The LAH correlations for each study group had a similar response to temperature with negative or no female and male correlations and positive or no female x male correlations (Table 6).

#### Yolk-sac Volume

Yolk-sac volume was highest for all study groups at 9.0°C and decreased as temperature decreased (Fig. 5). For cisco, the difference in YSV was similar between populations at the warmest incubation temperature (0.04 mm^3^) but diverged as incubation temperature decreased; YSV in LO-Cisco (0.40 mm^3^) was smaller than LS-Cisco (0.64 mm^3^) at the coldest temperature (Fig. 5). Yolk-sac volume in LS-Cisco and LO-Cisco responded differently to incubation temperature, with a 203.6 and 311.0% respective increase from the coldest to warmest treatment. For Lake Southern Konnevesi, the effect of temperature depended on species because the difference in YSV between LK-Vendace and LK-Whitefish was less pronounced at the coldest temperature (0.20 mm^3^) than at the warmest temperature (1.07 mm^3^; Fig. 5). LK-Vendace and LK-Whitefish had the strongest response to temperature with an increase in YSV of 445.0 and 536.6% from the coldest to warmest treatment, respectively. LK-Vendace and LK-Whitefish had a stronger, increasing response to increasing incubation temperatures in YSV than LS-Cisco and LO-Cisco (Fig. 5).

In the phenotypic variance component analysis, the residual error was the largest component of phenotypic variation in YSV (means >53.9%) for all study groups (Fig. 4, Online Resource 2). The mean YSV female variance was the highest percentage, excluding error, of the phenotypic variation for LK-Vendace (23.9%), LS-Cisco (20.5%), and LO-Cisco (23.9%; Fig. 4, Online Resource 2). The YSV correlations for female, female x male, and error variance components had differential responses to temperature with positive female, positive female x male, and negative error correlations for LK-Vendace, while LS-Ciso had inverse correlations to LK- Vendace (Table 6). All LO-Cisco variance components had no correlation to temperature (Table 6).

## DISCUSSION

Our incubation experiments demonstrated both similar and dissimilar reaction norms to temperature for developmental and morphological traits in coregonines. First, we found different responses to temperature in embryo survival within and among study groups. Second, incubation periods (both DPF and ADD) responded similarly to increasing temperature (negative response for DPF and positive response for ADD) among study groups, however, LK-Vendace had the strongest response and longest incubations across all temperatures. Third, all study groups had similar negative responses to temperature for LAH and positive responses in YSV, with the strongest responses for LAH in LS-Cisco and LO-Cisco and for YSV in LK-Vendace and LK- Whitefish. Lastly, differential levels of parental effects were found within and among study groups and traits.

Embryo survival had an overall negative correlation with increasing temperature among all study groups. However, cisco and vendace had different levels of response to temperature compared to European whitefish. LK-Whitefish embryo survival had the strongest, negative response to temperature (74.4% survival loss) and all other study groups were impacted less (< 26% survival loss) by increasing temperatures. Temperature is a strong driver of coregonine embryo development (Karjalainen et al., 2015) and survival (Colby & Brooke, 1970; Brooke & Colby, 1980; Luczynski & Kirklewska, 1984) but study groups showed different levels of sensitivity to increased incubation temperatures reflecting specific physiological adaptations.

Additionally, our experiment, temperature aside, provided near-optimal incubation conditions to individually reared embryos and these conditions are idealized compared to what occurs in the wild. For instance, embryos in the wild are deposited on the substratum and are exposed to deposited sediment that can impact survival (Müller, 1992). Interaction between temperature and sediments are likely, and temperature increases may act as a catalyzer of embryo sensitivity to sediment stress (Mari et al., 2016, 2021). Even though temperature did negatively impact embryo survival in our experiment, the effect of temperature in the wild, in combination with other factors, could be even stronger. Similarly, both study groups from North America required transportation, which could have had negative effects on embryo survival. Lake Superior had a greater transportation distance and time than Lake Ontario, which could explain some of the difference observed in embryo survival between the two study groups. Lake Ontario embryos had > 99% survival at optimal temperatures which suggests that transportation did not impact them.

The longer incubation periods from LK-Vendace and LK-Whitefish and strong response to increasing temperatures, even at the warmest incubation temperatures, support previous findings that vendace and European whitefish from Lake Southern Konnevesi have a high degree of developmental flexibility (Karjalainen et al., 2015). The different response between vendace and European whitefish was likely due to species-specific differences (Karjalainen et al., 2015) and ecotypes (*i.e.,* benthic versus pelagic; Mcphee et al., 2012). Additionally, the different magnitude of temperature responses among all study groups suggests a differential level of developmental plasticity to increasing incubation temperatures among species and locations. Long, relatively cold incubations may require a shorter period of spring warming for individuals to initiate hatching, while short, relatively warm incubations may require a longer period of warmer spring conditions to hatch (Karjalainen et al., 2015). If winter water temperatures rise as embryos incubate, the ability to match optimal spring nursery conditions may be weakened (Cushing, 1990; Karjalainen et al., 2015; Myers et al., 2015). Populations that are more resilient to increasing or variable winter incubation temperatures may have a better opportunity to regulate ontogeny and adjust the time of hatching.

Fish spawning strategies are variable, ranging in frequency from daily to once in a lifetime and in timing from the same time each year to across all seasons (McBride et al., 2015). For many species, spawning strategies and breeding patterns are constrained by the adult body condition, gonadogenesis, and the environment (Jørgensen et al., 2006; van Damme et al., 2009; Muir et al., 2014; McBride et al., 2015). In this context, the short duration of cisco embryo incubation periods when exposed to high temperatures was notable. High-latitude populations typically spawn earlier in autumn and may have the opportunity to delay spawning later into the season, while still providing an adequate incubation period for embryo development, if water temperatures continue to rise and do not inhibit ovulation as a result of climate change. However, low-latitude coregonine populations already spawn in late-autumn and early-winter (Stockwell et al., 2009; Eshenroder et al., 2016), which begs the question: do low-latitude populations have capacity to spawn later in the winter if temperatures continue to rise? Winter spawning may lead to less vulnerability to contemporary climate change. For instance, Atlantic herring (*Clupea harengus*) have both autumn- and winter-spawning stocks in the North Sea that share the same summer feeding grounds and start oocyte development at the same time (van Damme et al., 2009). Similarly, coregonines can exhibit varying spawning strategies.

Sympatric coregonine species with autumn, winter, and spring-spawning stocks co-occur in several northern- and central-Eurasian lakes (Eronen & Lahti, 1988; Schulz & Freyhof, 2003; Schulz et al., 2006; Ohlberger et al., 2008) and allopatric spring-spawning stocks of cisco are found in Lac des Écorces (southwestern Quebec; Hénault & Fortin, 1989, 1991; Pariseau et al., 1983). Winter and spring spawners continue oocyte development through autumn which results in a lower number of larger eggs compared to the autumn-spawning stocks (Eronen & Lahti, 1988; Hénault & Fortin, 1991). Oocyte development is driven by body energy content, and winter- and spring-spawning stocks may give iteroparous females the chance to mitigate the disproportionate energy demand toward somatic growth during the summer when metabolic demands are higher. Consequently, changes in the environment and the condition of an individual spawning adult could affect future coregonine spawning strategies. Our results suggest that the cisco embryos examined may not have the developmental plasticity to mitigate the effects of increased water temperatures on incubation period. In this context, adjusting the time of spawning may be a more efficient long-term life-history strategy than the embryos adapting to increased temperatures. Research on the reproductive plasticity of coregonine adults *(e.g.,* changes to fecundity of females and size of eggs from temperature) in the face of climate change is a logical next step.

Lake morphology is also important to consider for the question of a winter- or spring-spawning adaptation; deeper lakes could sufficiently provide cold thermal refuges at greater depths if suitable spawning habitat is available and level of oxygen sufficient (Jane et al., 2020). Spring-spawning ciscos in Lac des Écorces, where a 4°C summer stratum does not exist, initiate spawning when spring water temperatures reach 6°C at depths ranging from 20-30 m (Hénault & Fortin, 1989, 1991). This strategy of spawning in deeper, colder water allows for normal embryogenesis throughout the summer to mitigate high water temperatures during the summer period at shallow depths. Model projections of suitable thermal and oxythermal habitat for cisco indicate deeper and less eutrophic lakes will likely provide the best cold-water habitat as water temperatures and land uses change (Jacobson et al., 2010; Herb et al., 2014; Schmitt et al., 2020). While deep lakes may possess acceptable thermal refugia for coregonines, access to and requirements for suitable spawning and incubation habitat is unknown for most populations.

In addition to lower survival and shorter incubations as temperature increases, we also found both similar and different responses to temperature in morphological traits (*i.e.,* length-at-hatch and yolk-sac volume) among study groups. The dissimilarity in morphological traits among study groups is likely related to different egg sizes at fertilization. Smaller eggs will produce smaller larvae, requiring a lower growth and development rate and less demand on maternal yolk than larger eggs. The demand for yolk and egg size are positively related and temperature during embryogenesis is positively related to metabolic rate (Hodson & Blunt, 1986; Kamler, 2008). In all groups, LAH slightly decreases with increasing temperatures. In LK-Vendace and LK- Whitefish, however, YSV increases more strongly with temperature, than in cisco. This suggests a higher yolk conversion efficiency across all temperatures in LK-Vendace and LK-Whitefish than in cisco. All study groups had a decrease in YSV with time as basal metabolism consumed yolk as a function of the length of incubation. Regardless of the mode, our results suggest a synergistic relationship among species, location, egg size, incubation period, and incubation temperature in determining the phenotype of LAH and YSV.

The trade-off between LAH and YSV is well documented in larval fish physiology (Blaxter, 1991). Climate change impacts may only exacerbate the importance of each morphological trait in determining either a match or mismatch between larval coregonines and their zooplankton prey. While our experiments used constant incubation temperatures due to logistical constraints, the impact different spring warming rates can have on the time of hatching and the size of larvae should not be ignored. Lake Southern Konnevesi vendace and European whitefish previously exhibited flexibility in embryo development rates and feeding windows under different warming scenarios (Karjalainen et al., 2015). Such complex responses challenge our ability to predict the downstream impacts changing autumn, winter, and spring water temperatures may have on embryo and larval phenotypic plasticity.

Traits of embryos depend not only on species, population, and incubation temperature but also on parental and transgenerational effects (Blaxter, 1963, 1991; Kekäläinen et al., 2018). Our results suggest that both female and male effects controlled a portion of early-stage offspring trait phenotypes in coregonines. The variability in phenotypes induced by parental effects can provide more flexibility for a population to cope with changing inter-annual environmental conditions, prevent full year-class failure, and ensure population persistence (Wright & Trippel, 2009; Oomen & Hutchings, 2015; Karjalainen et al., 2016a). In fishes, the female effect is usually more pronounced than male and female x male interaction effects, and is stronger in traits directly related to egg size (Nagler et al., 2000; Kennedy et al., 2007; Huuskonen et al., 2011), and our results support this conclusion. Residual error estimates, however, remained high. Intersexual selection and mate pairing has been proposed as an important component affecting coregonine offspring fitness (Wedekind et al., 2008; Huuskonen et al., 2011; Karjalainen & Marjomäki, 2018), and may play a role in conserving natural biodiversity within populations (Anneville et al., 2015). The long-term stability of commercially exploited stocks, which can experience fisheries-induced evolution, has been linked to population diversity (Schindler et al., 2010; Freshwater et al., 2019). Spawning stocks that comprise individuals of variable sizes and ages (*e.g.,* portfolio effect; D. E. Schindler et al., 2010) may contribute differently to spawning, offspring performance, and recruitment (Luck et al., 2003; Figge, 2004), and are likely an important consideration as the rapid rate of climate change adds additional stressors on populations.

The methods we used provide reproducible experimental conditions (*e.g.,* uniform water source between laboratories, no moving water, minimal embryo disturbance, etc.) and standardized results that can be compared to future experiments that examine temperature responses across a wider range of populations. Additionally, efforts to include more northerly populations of cisco from Canada were thwarted due to restrictions on transport of live embryos across an international border. This further highlights the importance of the standardized experimental methods we used to allow for future large-scale, cross-laboratory experiments. However, our results do suggest that some form of latitudinal variation is likely present and promote fruitful opportunities for future large-scale experimental research on coregonines and other cold, stenothermic fishes.

Additionally, interpreting the impacts of parental responses within an environmental context continues to be important to determine how parental effects may mitigate species’ responses to rapid climate change. The existence of varying parental responses raises questions concerning possible causal mechanisms. Genomic studies will be needed to better understand what is genetically impacted by increasing temperatures, how it is impacted, and when during development (*i.e.,* when is temperature most critical; Chen et al., 2018; Narum et al., 2013). A mechanistic understanding of thermal response from populations across latitudes will be essential to predict the vulnerability of species and populations to climate change. Furthermore, incorporating phylogenetic contrasts would highlight the shared phylogenetic history and genetic architecture among our study groups and aid future comparative studies examining phenotypic traits across species and genera (Garland et al., 2005). This study does contain species from the same genus, but species are either reproductively isolated on different continents (*i.e.,* cisco and vendace) or are distinct ecotypes (*i.e.,* vendace and European whitefish). Moreover, stocking practices are important to consider, including supportive breeding, and may affect the adaptive potential of populations through an artificial selection process (Ford, 2002; Christie et al., 2012; Anneville et al., 2015) and their ability to respond to environmental changes.

## CONCLUSION

Water temperature is fundamental in regulating fish physiology, and environmental variation during development can play a large role in generating variability in offspring through phenotypic plasticity (Little et al., 2020). How coregonines respond during the critical embryonic stage is important to determine whether the capacity to respond to climate change and the projected increases in their incubation temperatures exists. Knowing how populations have adapted historically to environmental variability will help us understand the range of possible responses to climate change and assist managers to keep coregonines out of hot water.

## ACKNOWLEDGMENTS

We thank the staff at the Wisconsin Department of Natural Resources Bayfield Fisheries Field Station, United States Geological Survey (USGS) Tunison Laboratory of Aquatic Science, New York State Department of Environmental Conservation Cape Vincent Fisheries Station, and Konnevesi Research Station and local fishers for conducting field collections of spawning adults. We also thank Rachel Taylor, Mark Vinson, Daniel Yule, Caroline Rosinski, Jonna Kuha, and Rosanna Sjövik for help with fertilizations and experiment maintenance. This work was funded by the USGS under Grant/Cooperative Agreement No. G16AP00087 to the Vermont Water Resources and Lakes Studies Center and G17AC00042 to the University of Vermont. This work was made possible with funds made available to Lake Champlain by Senator Patrick Leahy through the Great Lakes Fishery Commission. We acknowledge INRAE, French National Research Institute for Agriculture, Food, and Environment, the UMR CARRTEL (INRAE - USMB) and the National Science Foundation (award number 1829451) for supporting a workshop to develop this experiment.

