## Supplementary material for "Influence of warming temperatures on coregonine embryogenesis within and among species": SI Table 1

SI Table 1. Phenotypic variance component analysis for embryo survival (%) and incubation period (number of days post-fertilization (DPF) and accumulated degree days (°C; ADD)) from Lake Southern Konnevesi vendace (LK-Vendace (*Coregonus albula*)), Lake Superior cisco (LS-Cisco (*C. artedi*)), and Lake Ontario cisco (LO-Cisco) across each incubation temperature treatment (°C).

| Trait | Study Group | T°C | Dam | | |  | Sire | | |  | Dam:Sire | | |  | Error | |
| --- | --- | --- | --- | --- | --- | --- | --- | --- | --- | --- | --- | --- | --- | --- | --- | --- |
|  |  |  | σ^2^ | *P* | % |  | σ^2^ | *P* | % |  | σ^2^ | *P* | % |  | σ^2^ | % |
| Embryo Survival | LK-Vendace | 2.2 | 1.19 | <0.001 | 24.53 |  | 0.12 | 0.481 | 2.53 |  | 0.24 | 0.211 | 4.92 |  | 3.29 | 68.03 |
|  |  | 4.0 | 0.68 | 0.004 | 16.41 |  | <0.01 | 0.999 | <0.01 |  | 0.16 | 0.064 | 3.97 |  | 3.29 | 79.62 |
|  |  | 6.9 | 0.89 | <0.001 | 20.25 |  | 0.20 | 0.159 | 4.57 |  | 0.03 | 0.799 | 0.70 |  | 3.29 | 74.48 |
|  |  | 8.0 | 0.33 | 0.007 | 8.33 |  | 0.20 | 0.074 | 4.97 |  | 0.14 | 0.078 | 3.49 |  | 3.29 | 83.21 |
|  | LS-Cisco | 2.0 | 1.61 | <0.001 | 29.82 |  | 0.0 | 1.0 | 0.0 |  | 0.50 | <0.001 | 9.25 |  | 3.29 | 60.94 |
|  |  | 4.4 | 2.82 | 0.007 | 36.45 |  | 0.0 | 1.0 | 0.0 |  | 1.62 | <0.001 | 20.97 |  | 3.29 | 42.58 |
|  |  | 6.9 | 1.51 | 0.094 | 20.32 |  | 0.0 | 1.0 | 0.0 |  | 2.64 | <0.001 | 35.44 |  | 3.29 | 44.24 |
|  |  | 8.9 | 0.44 | 0.082 | 9.66 |  | <0.01 | 0.999 | <0.01 |  | 0.78 | <0.001 | 17.29 |  | 3.29 | 73.04 |
|  | LO-Cisco | 2.0 | 0.55 | 0.450 | 11.28 |  | 0.0 | 1.0 | 0.0 |  | 1.01 | 0.295 | 20.90 |  | 3.29 | 67.82 |
|  |  | 4.4 | 1.59 | 0.007 | 32.55 |  | <0.01 | 0.999 | <0.01 |  | 0.0 | 1.0 | <0.01 |  | 3.29 | 67.45 |
|  |  | 6.9 | 0.87 | 0.003 | 19.30 |  | 0.0 | 1.0 | 0.0 |  | 0.34 | 0.057 | 7.65 |  | 3.29 | 73.06 |
|  |  | 8.9 | 0.71 | <0.001 | 16.48 |  | 0.22 | 0.008 | 5.12 |  | 0.10 | <0.001 | 2.24 |  | 3.29 | 76.17 |
| Incubation Period (DPF) | LK-Vendace | 2.2 | 8.95 | <0.001 | 21.27 |  | 0.87 | 0.160 | 2.08 |  | 0.0 | 1.0 | 0.0 |  | 32.25 | 76.65 |
|  |  | 4.0 | 149.67 | <0.001 | 30.03 |  | 26.12 | 0.023 | 5.24 |  | 9.79 | 0.138 | 1.96 |  | 312.80 | 62.76 |
|  |  | 6.9 | 49.61 | <0.001 | 18.93 |  | 0.92 | 0.815 | 0.35 |  | 10.18 | 0.008 | 3.89 |  | 201.37 | 76.84 |
|  |  | 8.0 | 62.35 | <0.001 | 42.32 |  | 1.02 | 0.693 | 0.69 |  | 7.26 | 0.001 | 4.93 |  | 76.69 | 52.06 |
|  | LS-Cisco | 2.0 | 233.62 | <0.001 | 34.48 |  | 0.0 | 1.0 | 0.0 |  | 111.34 | <0.001 | 16.43 |  | 332.54 | 49.08 |
|  |  | 4.4 | 30.81 | <0.001 | 27.53 |  | 5.82 | 0.115 | 5.20 |  | 7.32 | <0.001 | 6.54 |  | 67.95 | 60.72 |
|  |  | 6.9 | 7.04 | 0.001 | 18.37 |  | 1.83 | 0.089 | 4.76 |  | 1.51 | 0.024 | 3.95 |  | 27.95 | 72.92 |
|  |  | 8.9 | 1.03 | 0.615 | 3.80 |  | 0.65 | 0.782 | 2.42 |  | 9.01 | <0.001 | 33.34 |  | 16.34 | 60.44 |
|  | LO-Cisco | 2.0 | 123.54 | <0.001 | 61.37 |  | 15.66 | <0.001 | 7.78 |  | 1.70 | 0.002 | 0.84 |  | 60.41 | 30.01 |
|  |  | 4.4 | 67.65 | <0.001 | 48.78 |  | 16.22 | <0.001 | 11.70 |  | 1.60 | 0.002 | 1.16 |  | 53.20 | 38.37 |
|  |  | 6.9 | 25.00 | <0.001 | 48.27 |  | 3.19 | <0.001 | 6.16 |  | 0.34 | 0.086 | 0.66 |  | 23.25 | 44.91 |
|  |  | 8.9 | 7.77 | <0.001 | 29.95 |  | 0.80 | 0.012 | 3.10 |  | 0.44 | 0.032 | 1.70 |  | 16.93 | 65.26 |
| Incubation Period (ADD) | LK-Vendace | 2.2 | 187.78 | <0.001 | 24.59 |  | 29.01 | 0.024 | 3.80 |  | 0.0 | 1.0 | 0.0 |  | 546.95 | 71.62 |
|  |  | 4.0 | 3,250.95 | <0.001 | 30.56 |  | 549.52 | 0.025 | 5.17 |  | 210.28 | 0.142 | 1.98 |  | 6,625.67 | 62.29 |
|  |  | 6.9 | 2,622.69 | <0.001 | 18.98 |  | 49.01 | 0.813 | 0.35 |  | 538.27 | 0.007 | 3.89 |  | 10,610.13 | 76.77 |
|  |  | 8.0 | 3,978.98 | <0.001 | 41.44 |  | 36.31 | 0.825 | 0.38 |  | 489.35 | 0.001 | 5.10 |  | 5,097.39 | 53.09 |
|  | LS-Cisco | 2.0 | 793.55 | <0.001 | 33.84 |  | 0.42 | 0.997 | 0.02 |  | 399.38 | <0.001 | 17.03 |  | 1,151.61 | 49.11 |
|  |  | 4.4 | 572.77 | <0.001 | 27.51 |  | 108.43 | 0.115 | 5.21 |  | 136.38 | <0.001 | 6.55 |  | 1,264.28 | 60.73 |
|  |  | 6.9 | 327.96 | <0.001 | 18.34 |  | 85.05 | 0.090 | 4.76 |  | 70.59 | 0.024 | 3.95 |  | 1,304.32 | 72.95 |
|  |  | 8.9 | 80.39 | 0.600 | 3.84 |  | 51.62 | 0.777 | 2.47 |  | 693.60 | <0.001 | 33.15 |  | 1,267.02 | 60.55 |
|  | LO-Cisco | 2.0 | 388.91 | <0.001 | 59.03 |  | 54.46 | <0.001 | 8.27 |  | 5.74 | 0.003 | 0.87 |  | 209.73 | 31.83 |
|  |  | 4.4 | 1,258.57 | <0.001 | 48.60 |  | 303.66 | <0.001 | 11.73 |  | 30.77 | 0.001 | 1.19 |  | 993.64 | 38.49 |
|  |  | 6.9 | 1,167.10 | <0.001 | 48.34 |  | 148.69 | <0.001 | 6.16 |  | 15.58 | 0.090 | 0.65 |  | 1,082.77 | 44.85 |
|  |  | 8.9 | 599.46 | <0.001 | 29.96 |  | 62.09 | 0.012 | 3.10 |  | 34.07 | 0.031 | 1.70 |  | 1,305.37 | 65.24 |
