## Supplementary material for "Influence of warming temperatures on coregonine embryogenesis within and among species": SI Table 2

SI Table 2. Phenotypic variance component analysis for length-at-hatch (mm) and yolk-sac volume (mm^3^) from Lake Southern Konnevesi vendace (LK-Vendace (*Coregonus albula*)), Lake Superior cisco (LS-Cisco (*C. artedi*)), and Lake Ontario cisco (LO-Cisco) across each incubation temperature treatment (°C).

| Trait | Study Group | T°C | Dam | | |  | Sire | | |  | Dam:Sire | | |  | Error |  |
| --- | --- | --- | --- | --- | --- | --- | --- | --- | --- | --- | --- | --- | --- | --- | --- | --- |
|  |  |  | σ^2^ | *P* | % |  | σ^2^ | *P* | % |  | σ^2^ | *P* | % |  | σ^2^ | % |
| Length-at-Hatch | LK-Vendace | 2.2 | 0.06 | <0.001 | 37.01 |  | 0.02 | 0.011 | 11.06 |  | 0.0 | 1.0 | 0.0 |  | 0.08 | 51.93 |
|  |  | 4.0 | 0.07 | <0.001 | 41.39 |  | 0.02 | 0.007 | 9.74 |  | 0.0 | 1.0 | 0.0 |  | 0.09 | 48.87 |
|  |  | 6.9 | 0.08 | <0.001 | 49.84 |  | 0.0 | 1.0 | 0.0 |  | 0.01 | 0.227 | 3.82 |  | 0.08 | 46.34 |
|  |  | 8.0 | 0.05 | <0.001 | 34.13 |  | 0.01 | 0.239 | 6.14 |  | 0.02 | 0.033 | 9.92 |  | 0.08 | 49.81 |
|  | LS-Cisco | 2.0 | 0.49 | <0.001 | 61.71 |  | 0.01 | 0.490 | 1.55 |  | 0.01 | 0.783 | 0.86 |  | 0.28 | 35.87 |
|  |  | 4.4 | 0.20 | <0.001 | 30.63 |  | <0.01 | 0.999 | <0.01 |  | <0.01 | 0.910 | 0.56 |  | 0.44 | 68.81 |
|  |  | 6.9 | 0.18 | 0.006 | 31.62 |  | 0.0 | 1.0 | 0.0 |  | 0.11 | 0.007 | 18.89 |  | 0.28 | 49.48 |
|  |  | 8.9 | 0.10 | 0.001 | 29.00 |  | 0.01 | 0.465 | 3.17 |  | 0.0 | 1.0 | 0.0 |  | 0.23 | 67.83 |
|  | LO-Cisco | 2.0 | 0.12 | <0.001 | 29.51 |  | <0.01 | 0.908 | 0.23 |  | 0.0 | 1.0 | 0.0 |  | 0.28 | 70.26 |
|  |  | 4.4 | 0.07 | <0.001 | 13.63 |  | <0.01 | 0.999 | <0.01 |  | 0.0 | 1.0 | 0.0 |  | 0.42 | 86.37 |
|  |  | 6.9 | 0.04 | 0.001 | 16.36 |  | 0.01 | 0.208 | 4.66 |  | 0.84 | 0.840 | 1.02 |  | 0.19 | 77.96 |
|  |  | 8.9 | 0.02 | 0.078 | 8.95 |  | 0.0 | 1.0 | 0.0 |  | 0.05 | 0.050 | 12.60 |  | 0.21 | 78.44 |
| Yolk-sac Volume | LK-Vendace | 2.2 | <0.01 | 0.007 | 14.06 |  | 0.0 | 1.0 | 0.0 |  | <0.01 | 0.830 | 1.12 |  | <0.01 | 84.82 |
|  |  | 4.0 | <0.01 | 0.005 | 20.51 |  | <0.01 | 0.179 | 6.71 |  | <0.01 | 0.373 | 4.69 |  | <0.01 | 66.38 |
|  |  | 6.9 | <0.01 | 0.007 | 28.09 |  | <0.01 | 0.991 | 0.07 |  | <0.01 | <0.001 | 23.51 |  | <0.01 | 43.19 |
|  |  | 8.0 | <0.01 | <0.001 | 32.74 |  | <0.01 | 0.697 | 1.39 |  | <0.01 | 0.008 | 9.53 |  | <0.01 | 35.93 |
|  | LS-Cisco | 2.0 | 0.02 | 0.030 | 25.48 |  | 0.0 | 1.0 | 0.0 |  | 0.01 | 0.001 | 20.11 |  | 0.04 | 54.42 |
|  |  | 4.4 | 0.01 | 0.008 | 33.64 |  | <0.01 | 0.784 | 1.78 |  | 0.01 | 0.050 | 13.49 |  | 0.02 | 51.10 |
|  |  | 6.9 | <0.01 | 0.151 | 14.07 |  | <0.01 | 0.846 | 1.81 |  | 0.01 | 0.058 | 17.15 |  | 0.04 | 66.98 |
|  |  | 8.9 | 0.02 | 0.230 | 8.87 |  | 0.01 | 0.487 | 4.93 |  | 0.02 | 0.192 | 10.90 |  | 0.13 | 75.31 |
|  | LO-Cisco | 2.0 | 0.02 | <0.001 | 39.32 |  | <0.01 | 0.816 | 0.39 |  | 0.0 | 1.0 | 0.0 |  | 0.03 | 57.63 |
|  |  | 4.4 | 0.02 | <0.001 | 56.43 |  | <0.01 | 0.468 | 1.57 |  | 0.01 | 0.293 | 2.89 |  | 0.01 | 39.11 |
|  |  | 6.9 | 0.04 | <0.001 | 41.84 |  | <0.01 | 0.999 | <0.01 |  | <0.01 | 0.096 | 5.55 |  | 0.05 | 52.61 |
|  |  | 8.9 | 0.05 | <0.001 | 32.90 |  | <0.01 | 0.699 | 0.80 |  | 0.0 | 1.0 | 0.0 |  | 0.10 | 66.30 |
